# Single-Cell Transcriptomic Analysis Reveals Molecular Diversity of Human Oligodendrocyte Progenitor Cells

**DOI:** 10.1101/2020.10.07.328971

**Authors:** Xitiz Chamling, Alyssa Kallman, Cindy Berlinicke, Prajwal Devkota, Joseph L. Mertz, Calvin Chang, Aniruddha Kaushik, Liben Chen, Peter A. Calabresi, Hai-Quan Mao, Tza-Huei Wang, Donald J. Zack

**Affiliations:** Department of Ophthalmology, Wilmer Eye Institute, Johns Hopkins University School of Medicine, Baltimore, MD 21287, USA; Institute of Genetic Medicine, Johns Hopkins University School of Medicine, Baltimore, MD 21287, USA; Department of Computer Science, University of Miami, Coral Gables, FL 33146, USA; Department of Biomedical Engineering, Johns Hopkins School of Medicine, Baltimore, MD 21205, USA; Department of Mechanical Engineering, Johns Hopkins University, Baltimore, MD 21218; Department of Neurology, Johns Hopkins University School of Medicine, Baltimore, MD 21287, USA; Translational Tissue Engineering Center, Johns Hopkins School of Medicine, Baltimore, MD 21287, USA; Institute for NanoBioTechnology, Johns Hopkins University, Whiting School of Engineering Baltimore, MD 21218, USA; The Solomon H. Snyder Department of Neuroscience, Johns Hopkins University School of Medicine, Baltimore, MD 21205, USA

## Abstract

Injury and loss of oligodendrocytes can cause demyelinating diseases such as multiple sclerosis. To improve our understanding of oligodendrocyte development, which could facilitate development of remyelination-based treatment strategies, we performed single-cell-transcriptomic-analysis of developing human oligodendrocyte-precursor-cells (hOPCs). We engineered knock-in hESC-reporter lines in which an Identification-and-Purification tag is expressed under control of the endogenous, OPC-specific, *PDGFRα* promoter, and performed time-course single-cell-RNA-sequencing of purified hOPCs. Our analysis uncovered marked transcriptional heterogeneity of PDGFR*α*+ hOPCs and identified regulatory genes and networks that control their differentiation and myelination competence. Pseudotime trajectory analysis revealed two distinct trajectories for the development of oligodendrocytes vs astrocytes from hOPCs. We also identified novel transcription factors and other genes that developing hOPCs potentially use to choose between oligodendrocyte vs astrocyte lineages. In addition, pathway enrichment analysis followed by pharmacological intervention of those pathways confirm that mTOR and cholesterol biosynthesis signaling pathways are involved in maturation of oligodendrocytes from hOPCs.

## INTRODUCTION

Myelin, the insulating material that coats and protects axons and enables rapid saltatory conduction, is essential for the health and function of neurons^1^. Myelin disorders, the most common of which is multiple sclerosis, can be inherited or acquired, can occur from diverse etiologies such as genetic mutation, toxic injury, or autoimmune insult, and often lead to severe disability^2^. Although there are a number of drugs that can modulate the demyelinating process, these drugs are generally not effective at promoting remyelination. Development of remyelination-based therapies, which could have enormous clinical impact, would be greatly aided by increased understanding of the regulatory pathways and molecular mechanisms involved in the development of oligodendrocytes (OLs), a subtype of glial cells that are responsible for synthesizing and maintaining central nervous system (CNS) myelin. Transcriptomic and regulatory pathway studies, which has led to discovery of compounds that potentially target myelinogenic oligodendrocytes^3–6^, have predominantly used murine primary or mouse ES-derived OPCs and OLs. Although few transcriptomic studies that use microarray analysis or bulk RNAseq on human oligodendrocyte lineage cells (OLLCs) have been reported, more detailed studies using human OLLCs remain to be done^7,8,9^.

One of the bottlenecks limiting the use of primary human OLLCs in developmental and transcriptomic studies is the challenge of obtaining sufficient numbers of cells - primary human OPCs are rare, difficult to isolate, and cannot be expanded following isolation^7^. An alternative to using primary cells is to use human pluripotent stem cell (hPSC)-derived OPCs, but tracking and isolating pure OPCs from a mixed population of differentiating CNS cells is still technically challenging^10–13^. In this study, we engineered a unique hOPC reporter system by knocking-in an identification-and-purification (IAP) tag at the endogenous *PDGFRα* locus of a human embryonic stem cell (hESC) line. This reporter system enables scalable differentiation and purification of hOPCs at various stages of OLLC differentiation. The hESC-derived, purified hOPCs were then used for droplet-based single-cell capture and RNA-sequencing (scRNA-seq) at three different stages of differentiation. The scRNAseq, which allows identification of transcriptionally distinct cells within a population, facilitated an in-depth developmental analysis^14,15^ and revealed the diversity and genetic complexity of human OPCs. Transcriptomic and pathway enrichment analyses followed by pharmacological testing of the implicated pathways validated in human OPCs a number of regulatory genes and pathways that had been previously identified from murine studies^3,6,16–22^. We also found that unlike murine OPCs that mature predominantly into oligodendrocytes^9,23^, a subset of PDGFR*α*+ hOPCs can also give rise to astrocytes, and that a fraction of PDGFR*α*+ cells express mature astrocytes or oligodendrocytes markers. Taking advantage of the bipotential nature of our reporter OPCs, we performed a pseudotime analysis to track their differentiation trajectories^24^. This analysis identified novel genetic factors that are enriched in OLs or astrocytes, and are potentially involved in regulating OL vs astrocyte lineage specification.

## RESULTS

### Generation of an OPC differentiation and purification stem cell reporter line

Although several protocols for differentiation of OLLCs from hPSCs have been developed, generally, OPCs make up only about 50% of the resulting differentiated cell population^10,25,26^. The OPCs in a population can be purified using antibodies against O4 or A2B5 surface antigens. However, the majority of the O4+ cells represent post-mitotic immature oligodendrocytes and the A2B5+ cells consist of a heterogeneous population of glial restricted cells and developing neurons^7^. Therefore, as a first step to establish an improved platform for the study of human OL development, we created an OPC reporter system based on PDGFR*∝* expression and the identification-and-purification paradigm that we previously developed for the purification and analysis of human retinal ganglion cells^27^. Since *PDGFRα* is a well-characterized marker for myelinogenic OPCs ^7,25^ we hypothesized that a PDGFRα reporter could be utilized to identify, track, and purify developing OPCs from hESC-derived CNS cultures.

The embryonic stem (ES) line WA09 was engineered to express an IAP tag under control of the endogenous *PDGFRα* promoter (Figure 1). The IAP tag consists of a tdTomato florescent marker and a mouse cell-surface protein, Thy1.2, separated from each other and from the endogenous *PDGFR*∝ gene product by the “ribosome-skipping” 2A peptide (P2A-tdTomato-P2A-Thy1.2) (Figure 1a). Upon *PDGFRα* expression, tdTomato is expressed and localized to the cytoplasm and Thy1.2 is localized to the cell surface, allowing PDGFRα expressing cells to be immunopurified via Thy1.2 antibody conjugated magnetic microbeads (Figure 2b)^27^. Of note, the antibodies against mouse Thy1.2 used in the purification are species-specific and do not react against human Thy1^27,28^. CRISPR-based genome editing, using SpCas9 and a single-guide RNA (sgRNA) sequence designed to target the last exon of the human *PDGFRα* gene to stimulate homology-directed repair with a donor construct encoding the IAP tag flanked by 1 kb homology arms, was used to target the IAP tag to the *PDGFRα* locus^29^ and generate the PDGFRα-P2A-tdTomato-P2A-Thy1.2 hES reporter clone (PD-TT) (Figure 1a-c).

**Figure 1.**
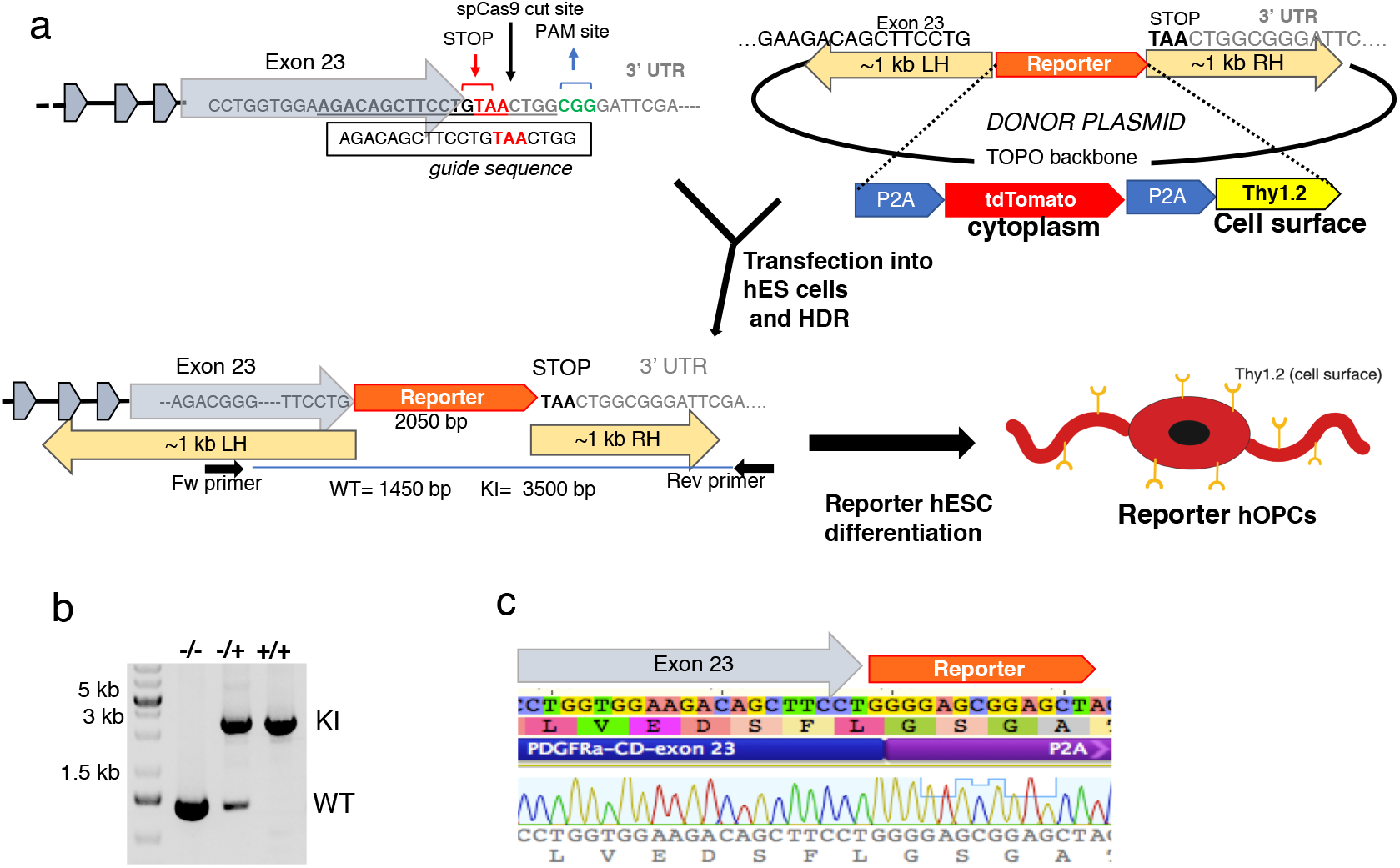
Generation of PDGFRα reporter ES cell line. **a)** Schematic diagram of IAP reporter knock-in into the *PDGFRα* locus using CRISPR-Cas9 genome editing. A plasmid containing spCas9 sequence and a guide sequence targeting the stop codon of the *PDGFRα* gene, and a donor plasmid containing reporter sequence flanked by 1kb homology arms was transfected into H9 ES cells. Following single cell passaging, PCR-based genotyping was performed on individual clones using a primer set that spans the reporter sequence (one primer at the homology arm and the other outside the homology arm), that allows differentiation between WT, heterozygous and homozygous knock-in clones **b)**. **c)** Sanger sequencing of the knock-in band to confirm insertion of the reporter at the correct location.

**Figure 2.**
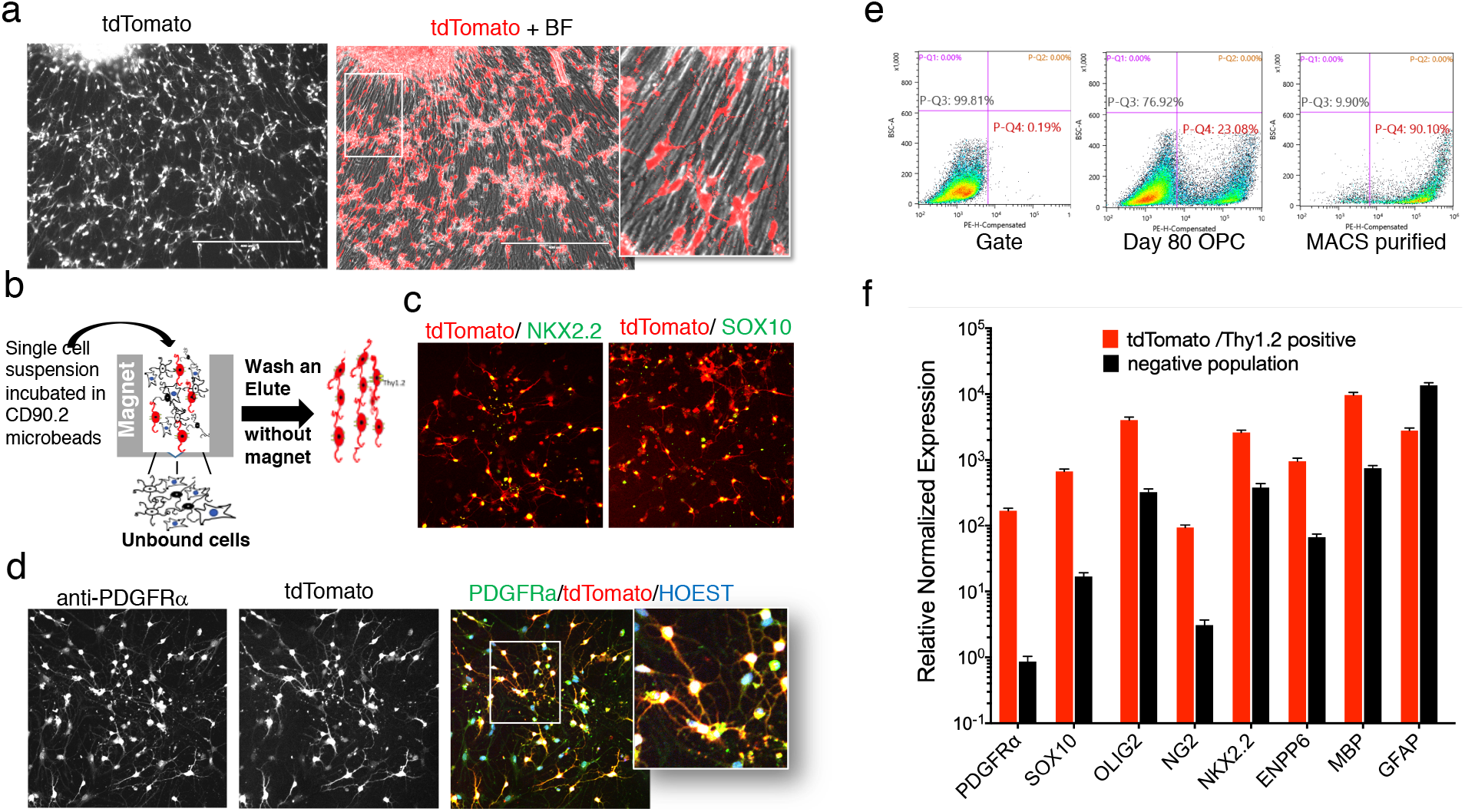
Differentiation and purification of PD-TT hESC reporter cell line. **a)** 80 days old OPC culture expressing tdTomato driven by the endogenous *PDGFRα* promoter. **b)** Schematic figure of MACS-based immunopurification of reporter cells using anti-Thy1.2 microbeads and a magnetic column. **c and d)** Immunohistochemistry demonstrating that the MACS purified cells co-express tdTomato and the OPC markers NKX2.2 and SOX10 **(c)** and PDGFRα **(d). e)** FACS analysis indicating 24.7% tdTomato+/PDGFRα+ cells at day 78, which is enriched to 90.10% after MACS purification. **f)** qPCR analysis shows enrichment of OPC markers in PDGFRα+/tdTomato+ cells compared to the tdTomato negative population.

### Differentiation and purification of PD-TT-derived OPCs

PDGFRα expressing OPCs initially emerge from the motor neuron progenitor domain (pMN) of the developing spinal cord^10,30^. Spinal cord patterning, pMN domain specification, and OPCs production have been modeled *in vitro* using human iPSCs^11,12,25^. Following neural patterning and neural stem cell differentiation via dual smad inhibition^31^, retinoic acid (RA) and a Sonic hedgehog (SHH) activator are added to promote OPC differentiation^12^. We used this differentiation strategy with the PD-TT reporter line (Figure S1a). Analogous to the timing of initial *PDGFRα* mRNA expression, small clusters of tdTomato+ cells were visible in our differentiation culture as early as day 8 (Figure S1b). However, morphologically bipolar, individual, tdTomato+ OPCs were not visible until approximately day 45, at which time mRNA levels of *PDGFRα* are increased ~700 fold compared to undifferentiated PD-TT cells (Figure S1c and Video S1). By day 60, numerous tdTomato+/PDGFR*α*+ cells were seen migrating out from neurospheres grown on poly-L-ornithine/laminin coated plates (Figure S1b and Video S2). By day 80, when grown in mitogen-free glial media, ~25% of the total cells in the differentiating cultures are tdTomato+ OPCs (Figure 2a and 2e). In addition, since the differentiated reporter OPCs also express the mouse Thy1.2 surface tag, these cells can be immunopurified via anti-Thy1.2 microbeads and magnetic activated cell sorting (MACS) (Figure 2b). Using MACS purification, we routinely obtain an approximately 90% pure population of tdTomato+ PDGFR*α* expressing cells (Figure 2e).

### PD-TT reporter cells express genes characteristic of OPCs and OLs

To further characterize the PDGFR*α/*tdTomato expressing cell populations, tdTomato+ and tdTomato-cell populations from day 80 differentiated cultures were obtained by MACS purification and the expression of various markers was analyzed by immunostaining and quantitative reverse-transcription PCR (qPCR) (Figures 2c, d, and f). The qPCR expression data were normalized relative to PD-TT undifferentiated stem cells. In the tdTomato+ cells, there was significantly enhanced expression of *PDGFRα* and other OPC markers, including *SOX10, OLIG2, CSPG4* (*NG2*), and *ENPP6. GFAP*, a gene whose express is highly enriched in astrocytes, was partially enriched in the tdTomato-population compared to the tdTomato+ cell population. Importantly, tdTomato+ cells expressed the Myelin Basic Protein (*MBP*) at ~10,000-fold higher levels compared to undifferentiated stem cells, and 13-fold over the tdTomato-cells (Figure 2f). The purified tdTomato+ cells differentiate into MBP+ oligodendrocytes in a mitogen free media (Figure 3a) within three weeks. When these cells are cultured on electrospun nanofibers, they align their processes along the nanofibers, and appear to myelinate them after three weeks in culture (Figure 3b). In addition, purified PDGFRα+/tdTomato+ cells maintain competence and capacity to mature into MBP+ OLs even after cryopreservation (Figure S2b and c).

**Figure 3.**
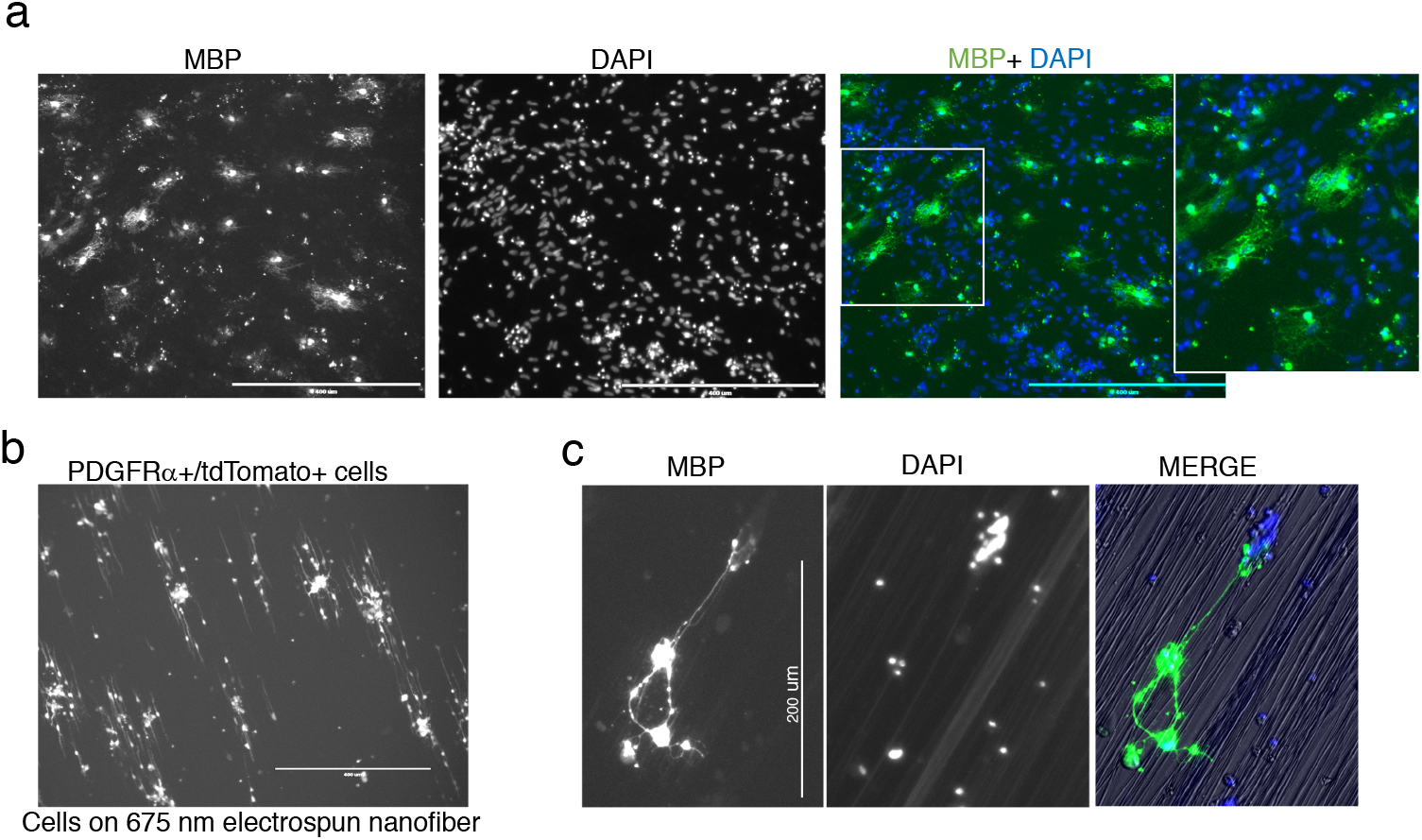
Maturation of reporter OPCs into myelinating oligodendrocytes. **a and b)** Day 80 MACS purified PDGFRα+/tdTomato+ cells were further cultured on PLO-laminin coated plates **(a)** or electrospun nanofibers of 675 nm diameter **(b)**. Cultures were grown in mitogen free media for 3 weeks and immunostained with MBP antibody. **b and c)** Within several days of culture, OPCs align their processes along the nanofibers.

### Single Cell Transcriptome analysis of OPCs

With the recent advent of technologies that allow scRNA-seq^14,15,32^, it is possible to simultaneously profile thousands of single cells and identify transcriptionally distinct cells within a population. To better understand the gene expression nuances associated with human OPC/OL differentiation, we applied the microfluidic based “Drop-seq” strategy^14^ to capture the transcriptome of differentiating OPCs at the singlecell level. Briefly, PD-TT reporter cells were differentiated for 77, 89 and 104 days; the samples were separately MACS purified (~90% purity), and single cells were captured with barcoded beads in droplets. cDNA and libraries for each sample were prepared independently, and an equimolar amount of each library was then pooled together for sequencing. From the three independent timepoints, a combined total of ~4,800 purified cells were captured. In order to eliminate probable doublets from the dataset, we bioinformatically filtered out cells with >30,000 unique molecular identifier (UMIs) (Figure S3a). Additionally, we removed cells that exhibited 1) expression of less than 250 genes or 2) greater than 20% mitochondrial gene content^33^. The remaining 3271 cells were used for further bioinformatic analysis.

T-distributed stochastic neighbor embedding (t-SNE)-based unbiased clustering of the 3271 cells clustered the tdTomato+/PDGFRα+ cells into 12 distinct populations (Figure 4a). To distinguish the cell-types represented by each cluster, we performed a spearman correlation of our dataset with a previously published RNAseq dataset from primary mouse and human CNS cells^3,34^ (Figure 4c and S4a). The clusters in our t-SNE plot share similarities with OPCs, early oligodendrocytes, mature oligodendrocytes, fetal astrocytes, and mature astrocytes. The cells that are grouped together in clusters 0, 1, and 3 at the center of the t-SNE plot represent precursor cells. More mature cells branch out to form more discrete clusters and seem to represent astrocytes lineage cells (ALCs) (clusters 5, 9 and 2) or oligodendrocyte lineage cells (OLLCs) (clusters 4, 7 and 8) (Figure 4, and Table 1). A relatively isolated cluster, cluster 10, shows enriched expression of numerous collagen-related (*COL1A1, COL1A2, COL3A1*)^35^ and actin-related (*ATAC2*) genes, and *PDGFRβ*^36^, which are known to be expressed in pericytes. Interestingly, some endothelial (*ANXA2*) and many mature astrocyte (*SLC40A1, IGFBP7, SPARC*) markers were also enriched in this cluster (Table 1). Except in one or two cells, expression of OPC markers was not detected in this cluster (Figure 4d).

**Figure 4.**
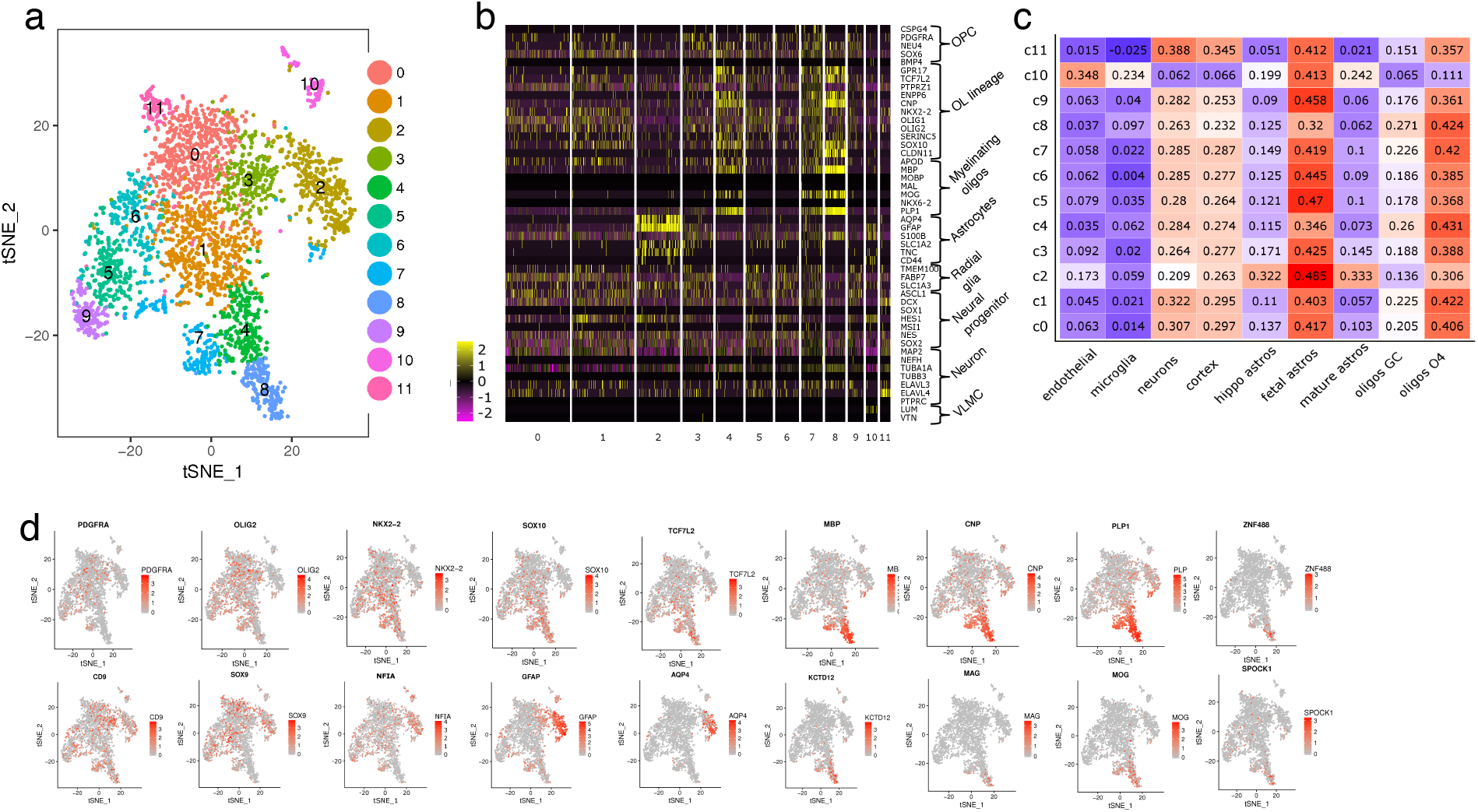
Single cell transcriptomic analysis of purified OPCs. **a)** t-SNE based unsupervised clustering of the single cell transcriptional profiles of purified PDGFRα+/tdTomato+ OPCs. Cells (each dot represents a cell) are divided into 12 clusters (0 through 11). **c)** Heatmap showing expression patterns for each cluster of previously reported markers of various CNS cell types. Cluster specific enrichment of various OL and astrocyte genes is apparent. **c)** Spearman correlation of the single cell transcriptome-based clusters with RNAseq data from primary human brain cells ^34^. **d)** Distribution of various genetic markers among different clusters presented as an enrichment heatmap.

**Table 1.**
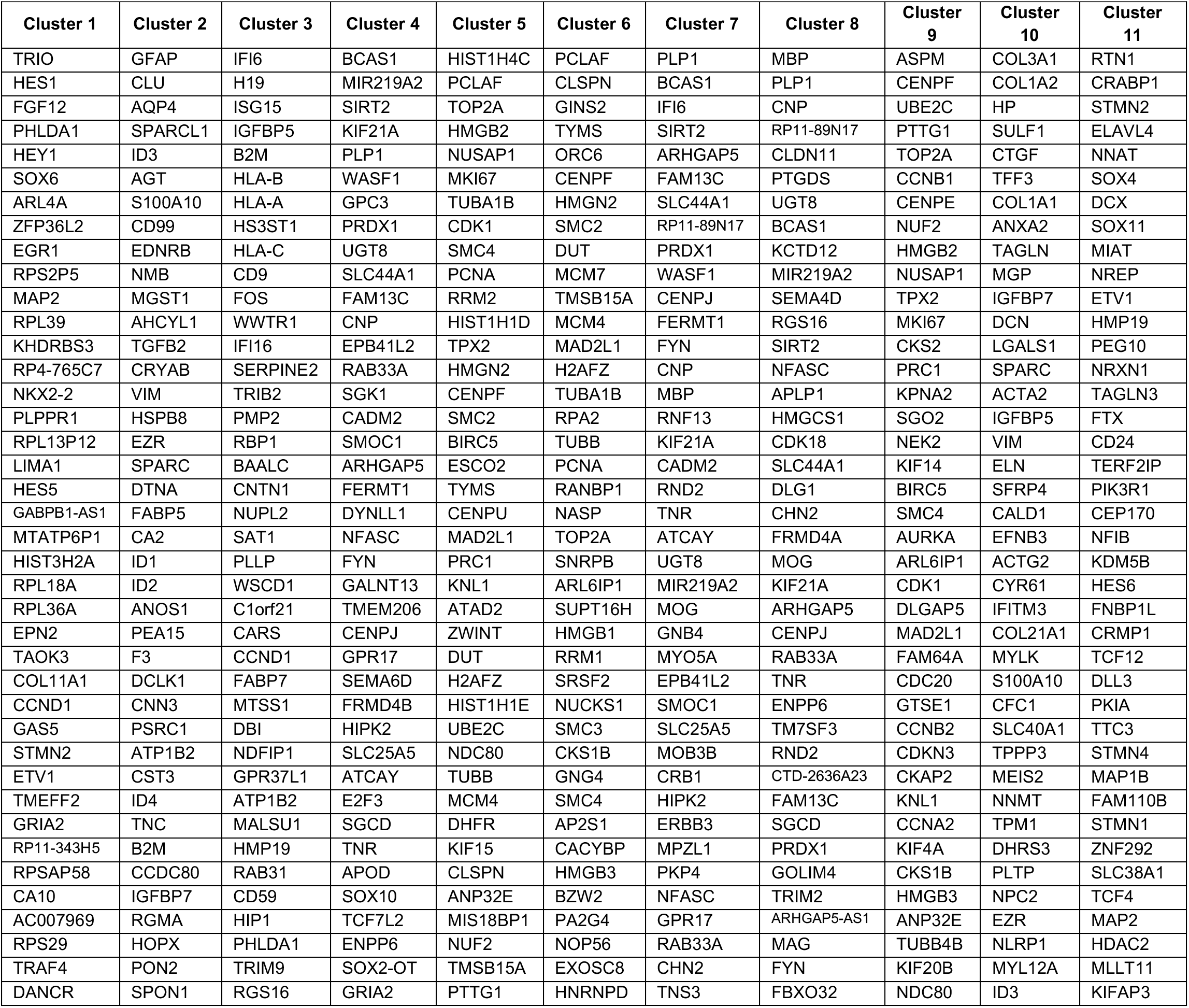
Top 40 differentially expressed genes per cluster ranked by fold change.

An expression heatmap of previously known markers for various CNS cell types also showed enrichment of OL genes in clusters 4, 7, and 8 and mature astrocyte-related genes in cluster 2 (Figure 4b)^7^. Mature OL markers such as *MAG, MOG*, and *ZNF488* were more enriched in cluster 8 (Figure 4d), indicating this as a more mature OL population than clusters 4 and 7, which are likely newly formed OLs (nfOLs). However, it should be noted that some previously described OPC (e.g. *BMP4*) and OL (e.g. *MOBP* and *MAL*) markers were not detected. Expression of markers for radial glia cells (*TMEM100, FABP7, SLC1A3*), which are known as progenitors of OLs, astrocytes and neurons in the CNS, were quite evenly distributed throughout the clusters (Figure 4b).

Of all the tdTomato+/PDGFR*α*+ cells analyzed, 901 (27.5%) expressed *GFAP* and 733 (22.4%) expressed *MBP* markers representing ALCs and OLLCs, respectively, which supports a previous report that human OPCs have the potential to differentiate into both astrocytes and oligodendrocytes^7^. We also separately analyzed the tdTomato+/PDGFR*α*+ cells from days 77, 89, and 104. Cells from each time-point formed distinct clusters with OPC, OL and astrocyte enriched populations, which confirms that purified hESC-derived OPCs, even at the same differentiation time-point, are temporally heterogenous in that they contain cells at different stages of differentiation (Figure S3c).

### ScRNA-seq reveals distinct cell type-enriched genes, some of which appear to be species-specific

We next examined other highly enriched and differentially expressed genes from each of the clusters illustrated in the t-SNE plot. The majority of the differentially expressed genes from the OL, nfOLs, fetal astrocytes, and mature astrocyte clusters match previous reports^3,34^. For example, *MBP, PLP1, CNP, CLDN11, UGT8, BCAS,1 SIRT2, MOG, MAG, TNR, ENPP6*, and *CHN2* were enriched in the OL and nfOL clusters (4, 7, and 8); and *GFAP, CLU, AQP4, ID3, EDNRB, MGST1, AHCYL1*, *EZR, HSPB8, SPARC, DTNA*, and *FABP5* were enriched in the astrocyte and fetal astrocyte clusters (2, 5, and 9) (Table 1, Table S2 and http://zacklab.org/OPCs/). Genes that have been reported to be specifically enriched in human OLs and astrocytes but not expressed in mouse OLs or astrocytes, such as *APCDD1, HMGCS1, PMP2* and *WIF1^34,37^*, were also enriched in the respective clusters as expected. However, we noted several subtle differences. In contrast to previous suggestions that CD9 is specifically expressed in the myelinogenic OLLCs^7^, we found quite even distribution of CD9 in all the clusters (Figure 4d), implying that the PDGFR*α*+/CD9+ cells do not unambiguously define myelinogenic progenitor cells in our stem cell-derived human OPCs. In addition, we found OL-specific enrichment of several genes (e.g. *KCTD12, SLC7A14, HMGCS1, SPOCK1, FAM13C, FAM131C, TMEM206*, and *KIF21A*) in our system (Table 1) that contrast with previous publications. For example, *HMGCS1*, which shows OL-specific enrichment in our data set has been reported to not be expressed in mouse CNS cells^3^, although it was previously reported to be essential for cholesterol biosynthesis and myelin production in zebrafish CNS^38^. Among human CNS cells, *HMGCS1* was reported to be enriched in fetal astrocytes^34^. *KCTD12, TMEM206, FAM131C, FRMD4B* and *APCDD1* were all enriched in human OLs in our system, but in the mouse, based on bulk RNAseq of purified CNS cells, *KCTD12* and *TMEM206* are enriched in microglia; *FAM131C* is enriched in neurons; and *FRMD4B* and *APCDD1* expression is specific to endothelial cells^3^. Interestingly, in bulk RNAseq of purified human CNS cells, some of these genes were enriched in cell-types other than OLs: for example, *KCTD12* was highly enriched in microglia, *FAM131C* was slightly enriched in mature astrocytes and neurons, *APCDD1* was expressed more in mature astrocytes and endothelial cells than in oligodendrocytes, and *SPOCK1* was more enriched in neurons than OLs^3,34^.

As noted above, when compared to previously published bulk RNAseq data, we found sub-populations of the tdTomato+/PDGFRα+ cells with very high correlation to human astrocytes. Clusters 2, 5, and 9 had comparatively higher correlation (>0.45) with fetal astrocytes (Figure 4c). Cluster 2, in addition, had relatively strong correlation with mature astrocytes, and showed enrichment of mature markers such as *SPARCL1, AQP4*, and *IGFBP7, AGT*, and *EDNRB* (Table 1, Table S2)^34,39^. Clusters 5 and 9 had highly enriched expression of proliferative genes associated with fetal astrocytes, such as *TOP2A, HIST1H4C, PCLAF, HMGB2, NUSAP1, MKI67* and *TMSB15A*, which were not expressed in cluster 2 (Table 1, Table S2). This data further confirms that cluster 5 and 9 represent fetal astrocytes and cluster 2 are more similar to mature astrocytes. In our mature astrocyte cluster (cluster 2), we also found enrichment of a few genes such as *S100A10, CD99, and ID2* that were previously implicated in endothelial cells or microglia in population-based RNAseq studies^3,34^. The same studies also reported very specific expression of *CRYAB* and enrichment of *CA2* in OLs, but they were specific to astrocytes in this study.

We additionally identified a number of differentially expressed primary microRNAs (pri-miRNAs) in our scRNAseq data (Figure S3b). For example, miR219-A2 was enriched in OL clusters while miR100HG and miR99AHG were enriched in astrocyte clusters. These differentially expressed pri-mRNAs are potentially involved in regulating the fate of an OPC to become either an OL or an astrocyte.

### Pathway enrichment analysis reveals pathways associated with oligodendrocyte and astrocyte differentiation

In order to get further insight into the molecular pathways that define OPCs, OLLCs or ALCs subpopulations, we performed pathway enrichment analyses on differentially expressed genes from each cluster using Ingenuity Pathway Analysis (IPA) and the Kyoto Encyclopedia of Genes and Genomes (KEGG) database. As expected, the majority of clusters that are at closer proximity to each other in the t-SNE plot (Figure 4a) share stronger correlation between enriched IPA canonical pathways (Figures 5a and b). However, cluster 2 and cluster 7 do not exhibit the same strong correlation with other ALC or OLLC clusters (Figures 5b and b), which indicates differential pathways between fetal and mature astrocytes, and suggests that cluster 7 could be a distinct subpopulation within the OL lineage. Numerous pathways with P-values denoting significance were identified for each cluster (Figure 5b and supplemental table 3). In the mature OL cluster, CXCR4, Sphingosine-1-phosphate (S1P), and integrin signaling pathways (ISP), which are known to be important for OPC maturation, oligodendrocyte survival and myelination were upregulated^20,21,40^, and EIF2 and RhoGDI signaling were downregulated (Figure 5d). Of additional interest, mTOR signaling and cholesterol biosynthesis pathways (CBP), known to be involved in OL differentiation and myelination, were enriched in the mature OL cluster, and Cdc42 and caveolar-mediated endocytosis signaling, which are implicated in astrocyte functions, were enriched in the mature astrocyte cluster^16,18,41,42^ (Figure 5c).

**Figure 5.**
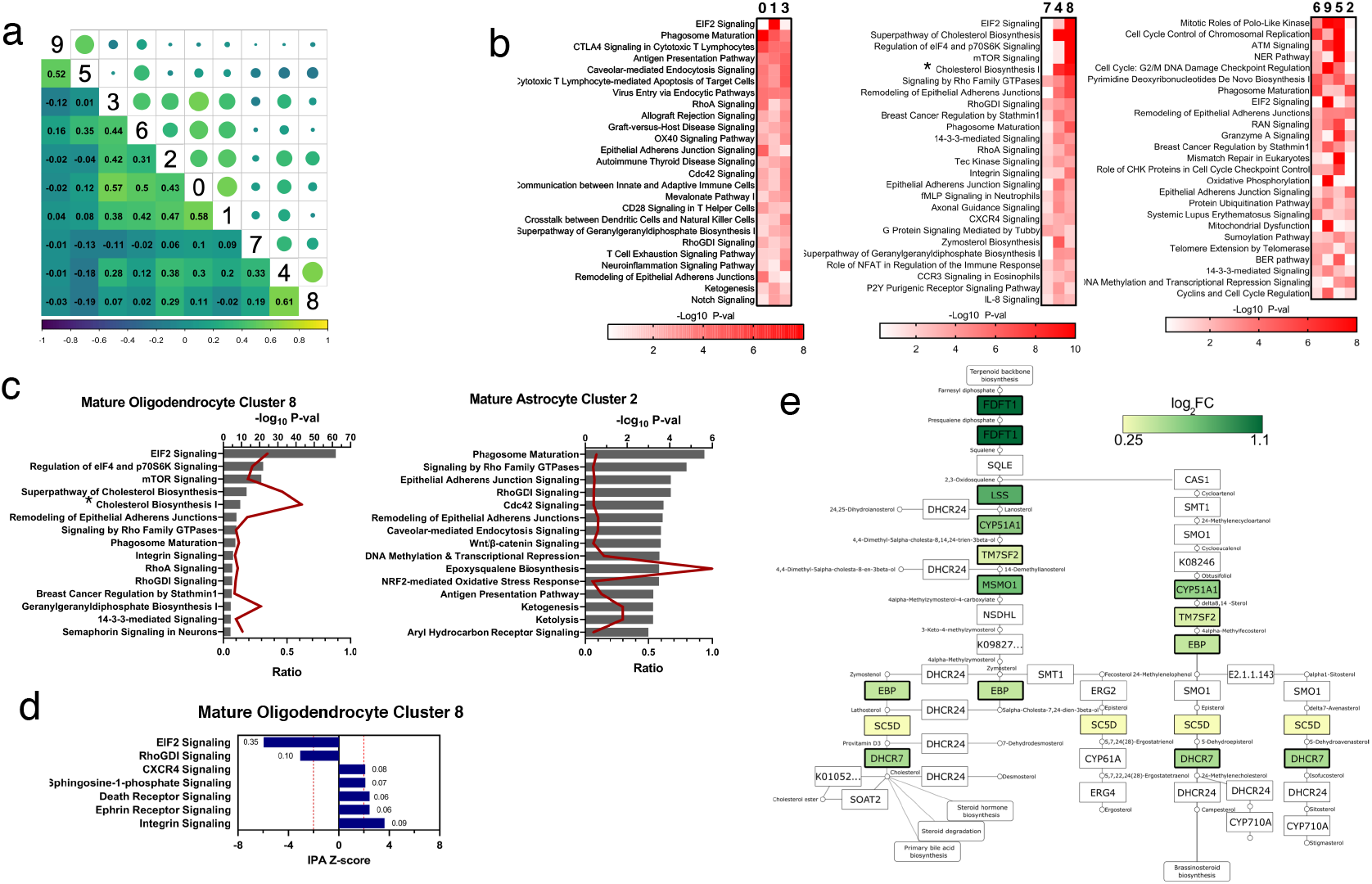
Pathway analysis of the differentially expressed genes. **a)** Pearson’s correlation of the pathways between each cluster. Circle diameters indicate strength and the colors provide directionality of the correlation. Actual values for each correlation are presented on the lower half of the graph. **b)** Top 25 pathways implicated for each cluster are presented as a heatmap. Numbers on the top represent the clusters, which are grouped together according to their predicted lineage: clusters 5, 6, 9, and 2 are astrocyte lineage; 4, 7, and 8 are oligodendrocyte lineage; and 0, 1, and 3 are OPCs. Colors indicate significance based on −log10 P-value. **c)** A combined bar and line chart for the pathways implicated in the mature astrocyte and the mature oligodendrocyte clusters. Top 15 pathways sorted based on their combined P-values are presented as a bar chart. The line chart represents overlap ratio (lower axis) of genes from scRNA-seq over the total number of genes predicted for that pathway. **d)** Pathways that are significantly differentially modulated in the mature OL cluster. Z-score for each pathway was calculated by IPA. The numbers next to each bar represent the overlap ratio. **e)** A map for KEGG steroid biosynthesis pathway (analogous to cholesterol pathway in IPA), which is highly enriched within the mature OL cluster (#8), is presented here as an example. Colored boxes denote genes that are significantly enriched based on our RNA-seq data. * represents all three cholesterol biosynthesis pathways (cholesterol biosynthesis I, II, and III).

To confirm the biological relevance of the above-described, bioinformatically implicated, pathways in hOPC to hOL maturation, we experimentally tested the consequences of pharmacologically inhibiting their activity in developing hOPCs. Ketoconazole, amorolfine and tasin-1, which target *CYP51A1, TM7SF2, and EBP* respectively, were used to target the cholesterol biosynthesis pathways; CYM5520 was used to target S1P; WZ811 was used to inhibit CXCR4 signaling; and rapamycin was used to inhibit mTOR signaling. MACS purified day 85 hOPCs were cultured in the presence of one of the compounds, or DMSO as control, for one week. The effects of the compounds were then tested by qPCR and immunostaining. Similar to a recent report^16^, inhibition of *CYP51A1, TM7SF2, and EBP* significantly increased MBP expression in our human system (Figures 6a and b). Inhibition of mTOR signaling by rapamycin significantly reduced the expression MBP and increased the expression of OPC markers, *PDGFRa* and *CSPG4* (*NG2*), which indicates that the mTOR signaling is essential for maturation of hOL from hOPCs. However, targeting S1P and CXCR4 pathways with our chosen compounds did not show any significant effect on hOL differentiation/maturation (Figure 6a).

**Figure 6.**
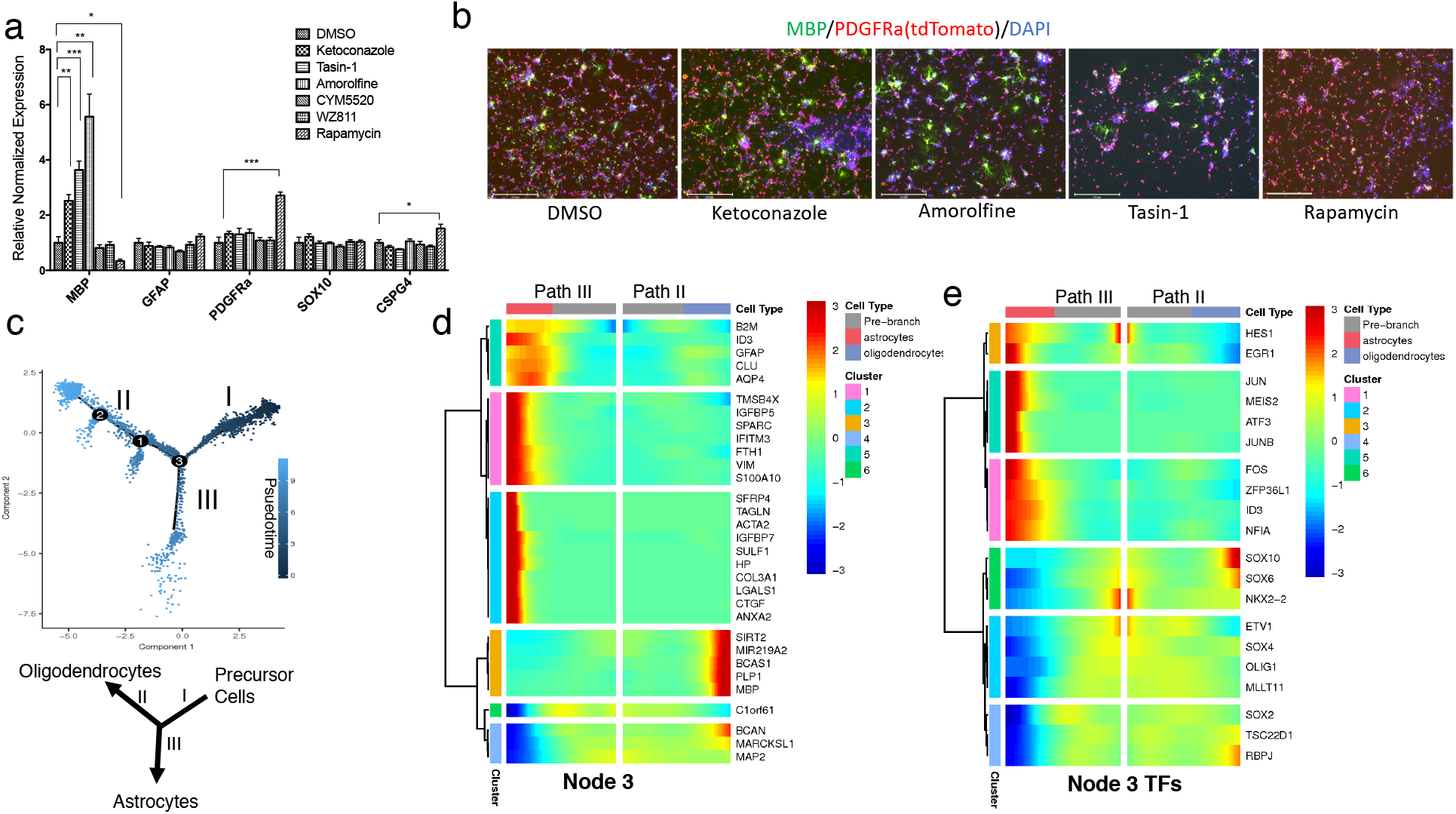
Effects of select small molecules on hOPCs and pseudotemporal trajectory analysis of the single cell transcriptomes of the hOPCs. **a and b)** Reporter hOPCs purified at day 85 of differentiation were cultured in the presence of compounds that inhibit CBP, S1P, CRCX4, or mTOR signaling pathways. The treated cells were tested by qPCR (a) and immunohistochemistry (b) to assess the effect on hOPC differentiation to OL - CBP inhibition promoted differentiation while mTOR inhibition had the opposite effect. **c**) Developmental trajectories of the reporter OPCs generated using Monocole-2 show two prominent paths for the OPCs. Darker colored dots (on Path I) represent developmentally younger cells and the lighter colored dots (on Paths II and III) represent more mature cells. **d and e)** Differential heatmap capturing the most differentially expressed genes **(d)** and transcription factors **(e)** between Paths II (oligodendrocytes) and III (astrocytes).

### Pseudotemporal trajectory analysis further defines the bi-potential nature of PDGFRα+ cells

We performed a monocle-based pseudotime analysis on our scRNAseq data to create a developmental trajectory tracing the lineage specification of PDGFRα+ OPCs as they mature. Analysis of the pseudotemporal trajectory presented two prominent paths for the precursor cells, indicating that the PDGFR*α*+ OPCs can follow two distinct cell lineages (Figure 6c). We examined the highly differentially expressed genes between the two trajectories and identified path II as OLLCs and path III as ALCs (Figure 6c-e). Among the differentially expressed genes, *BCAN, MARCKSL1*, and *SIRT2* were not only enriched in OLLCs but also significantly downregulated in astrocyte lineage cells, while the opposite was true for *B2M, TMSB4X*, and *FTH1*. To try to gain insight into the molecular mechanisms determining these distinct lineage pathways, we analyzed the transcription factors that are differentially expressed between the two paths. The TFs *HES1, EGR1, ZFP36L1*, and *NFIA*, and *JUNB* were enriched in ALCs but reduced in OLLCs, while expression of *SOX10, SOX4, SOX6, TSC22D1, and RBPJ* was enriched in OLLCs but reduced in ALCs (Figure 6e). Our data suggests that these transcription factors play a role in determining whether OPCs become OLs or astrocytes.

The monocle-based developmental trajectory analysis defined seven distinct cellular states (Figure 7a). Cells in state 1 were precursor cells, state 2 was comprised of astrocyte cells, and state 6 were oligodendrocyte cells. Along path II leading towards mature OLs, smaller branches consisting of state 5 and 7 cells diverged from the OL trajectory, and were interestingly enriched for astrocyte markers (*GFAP, AQP4, NFIA, SOX9*) rather than OL markers (*PLP, MBP, SOX10, TCF7L2, OLIG2, NKX2.2*) (Figure 7b and S5a). To further define the relationship between the cell states, we performed a correlation of the state 5 and 7 cells with state 2 (astrocytes) and state 6 (oligodendrocyte) cells using the average gene expression within each state. As expected, we found stronger correlation of cells in state 5 and 7 with astrocytes than oligodendrocyte cells (Figure 7c). We also overlaid the cells from each of the 12 clusters (Figure 4a) to the pseudotime trajectory and found that the cells from astrocyte cluster (cluster 2) make up the majority of state 5 and 7 population, and that the cells from OL clusters (cluster 4 and 7) make up the majority of the state 6 population (Figure S5c). These results suggest that astrocytes can also originate from PDGFR*α*+ cells at later stages of oligodendrocyte differentiation *in vitro*. Furthermore, lack of any divergent OL branch or OL state cells in the astrocyte branch (path III) indicates that emergence of oligodendrocytes from later stage astrocyte lineage cells is unlikely. Interestingly, the cells at the farthest end of astrocyte trajectory (path III) consisted of cells from cluster 10 (Figure S5c). This suggests that the cluster 10 cells, although expressed numerous pericyte marker genes, might be closer to a mature astrocyte population.

**Figure 7.**
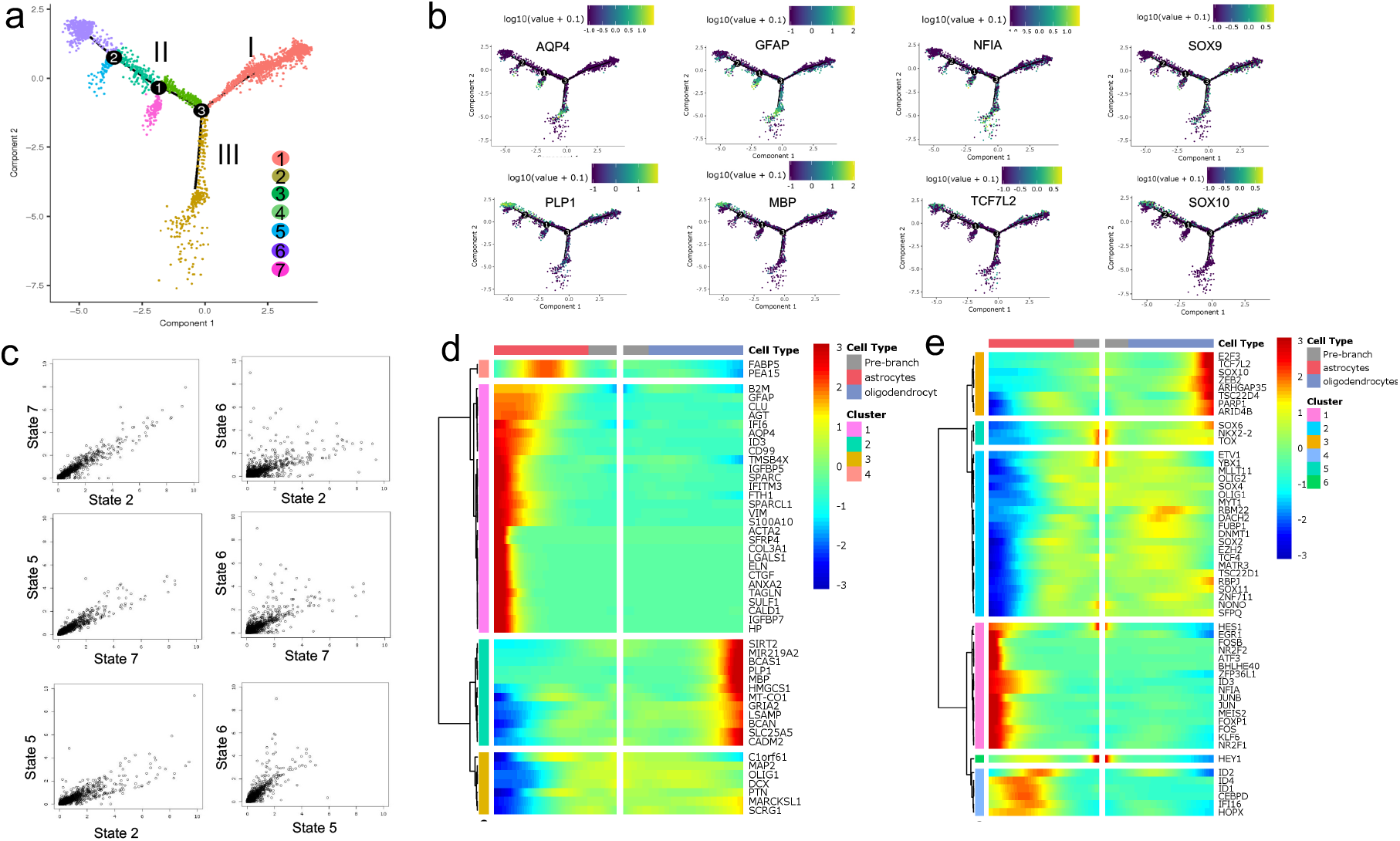
Pseudotemporal trajectory of differentiation based on single cell transcriptomes. **a)** Ordering cells along the trajectory divides the population into seven different states. State 1 cells are OPCs, state 2 cells are astrocytes, and state 6 cells are oligodendrocytes. **b)** Expression of OL and astrocyte markers within the trajectory further confirms Path III as the astrocyte and Path II as oligodendrocytes lineage cells. Astrocyte makers (*GFAP, AQP4, SOX9, NFIA*) are highly enriched in Path III and in the smaller branches that emerge from path II. OLLC markers (*SOX10, MBP, PLP, TCF7L2*) are enriched in Path II, but not in the smaller branches. **c)** Scatterplot showing the average gene expression of the cells in astrocyte vs oligodendrocyte states. Each dot represents a gene and axes are average number of transcripts per cell. Axes were limited to 10 in the plot in order to achieve better resolution. Cells from the small branches (state 5 and 7) have higher correlation with astrocytes (state 2 cells) than oligodendrocytes (state 6 cells). **d and e)** Heatmap of the most differentially expressed non-transcription factor **(d)** and transcription factor genes **(e)** between state 6 (oligodendrocyte cells) and state 2 (astrocyte cells).

Additionally, examination of the most differentially expressed genes and transcription factors at nodes 1 and 2, and in cells at states 2 vs state 6 (Figure 7d, e and S4), identified numerous differentially expressed genes of potential interest. In addition to TFs previously implicated in OPC/OL/astrocyte differentiation, such as *SOX10 and TCF7L2*, this analysis implicated a number of previously less characterized factors in OPC/OL/astrocyte differentiation, including *ZEB2, TSC22D4, ARID4B, PARP1, E2F3, and ARHGAP35*, which were significantly enriched in OLs, and *HES1, FOSB, NFIA, NR2F1*, and *ZFP36L1*, which were enriched in astrocyte cells. Numerous TFs such as *SOX4, SOX11, MLLT11, RBM22, ZNF711, EZH2*, and *DACH2* were slightly upregulated in OLs but highly downregulated in ALCs (Figure 7d).

Furthermore, each path of the trajectory consisted of cells from all three (day 77, 89, and 104) timepoints (Figure S5b), which further supports that PDGFR*α*+ cells are transcriptionally and developmentally heterogeneous, with some cells, even at the earliest timepoint tested (day 77), being already relatively mature.

## DISCUSSION

Recent advances in stem cell biology and differentiation methodology have led to the development of protocols for the generation of OPCs from hPSCs^10^, making possible detailed analysis of the molecular mechanisms underlying human OPC differentiation and myelination. One limitation of the current differentiation protocols, however, is that on average OPCs make up only about 50% of the resulting differentiated cell population. Here, in order to develop a simple and efficient method for obtaining highly purified populations of OPCs, we used CRISPR-based genome editing to introduce an IAP (P2A-tdTomato-P2A-thy1.2) tag into the endogenous *PDGFRα* locus. The resulting reporter cell line allows for optimization, scalable differentiation, and purification of human OLLCs (>90% PDGFR*α/*tdTomato-expressing cells) at different stages of differentiation. Unlike hPSC-derived OPCs that are purified using the O4 antigen^43^, OPCs purified by this method maintain high survival, competence, and the capacity to mature into MBP+ OLs even after long-term cryopreservation. This hOPC reporter/purification system provides a useful resource and a powerful tool for optimizing new, more efficient OPC differentiation protocols, enabling the easy quantification of effects that small molecules and patterning factors have on promoting OPC differentiation. In addition, the high capacity and simplicity of the methodology could aid in establishing a human OPC-based drug-discovery platform to screen for novel myelination promoting compounds.

To define the transcriptional diversity and heterogeneity of PDGFRα+ human OPCs, we performed scRNA-seq on the hPSC-derived reporter OPCs at various time points after the initiation of differentiation. Unbiased clustering of the cells identified 12 distinct clusters, the majority of which represented OPCs and cells committed to either the OL or astrocyte lineages. These findings are consistent with a previous report indicating that primary human OPCs can give rise to both astrocytes and OLs *in vitro*^7^. Although studies suggest that mouse OPCs exclusively generate OLs during normal development, A2B5+ rodent OPCs are capable of differentiating into type 2 astrocytes *in vitro* when cultured in the presence of serum or BMP4^44,45^. Given that ~27% of the PDGFRα+/tdTomato+ OPCs are also positive for A2B5 (Figure S2d), the astrocyte population presented in this study are likely to be type 2 astrocytes.

A variety of studies indicate that OPCs, but not differentiation-committed and mature OLs, express PDGFR*α*^1,4,10,46^. Thus, it was surprising that a significant number of our PDGFR*α*+/tdTomato+ cells showed strong correlation with OLs and astrocytes and expressed numerous mature markers (Figures 4–5, Table 1). For example, genes encoding synapse-inducing protein (*SPARCL1*) and transmembrane proteins (*CLU and SPON1*), which are expressed in mature astrocytes^34^, were highly enriched in the astrocyte cluster. Similarly, genes associated with myelin production (*MBP, MYCRF MAG, MOG*), which are expressed in mature OLs, were enriched in the OL cluster (Table 1 and Table S2). It should be noted, however, that the *PDGFRα* mRNA in our single cell data is not well detected in the clusters representing mature cells (Figure 4d). Possible explanations for why *PDGFRα* mRNA was detected in only a small fraction of the purified PDGFRα+/tdTomato+ cells, given that the tdTomato protein was obviously present at the time of purification, are that 1) *PDGFRα* mRNA in mature cells is transiently expressed and no longer present at the time of purification, or 2) perhaps, in the mature cells, PDGFR*α* mRNA is expressed as a low abundance transcript, and such transcripts are often missed in scRNAseq^14,15,32^. Nonetheless, our data suggests that the PDGFR*α*+ cells represent a significant pool of mature astrocytes and OLs, and that the expression of PDGFR*α* with other mature markers such as *PLP1, MBP, CNP, AQP4, SPARCL1, CLU* etc. is not mutually exclusive.

The identification of transcriptionally distinct cells within the PDGFR*α*+ population would not have been possible by standard bulk RNAseq. The single cell analysis identified several novel genes specific to human OLs (e.g. *KCTD12, SLC7A14, HMGCS1, SPOCK1, FAM13C, FAM131C, TMEM206*, and *KIF21A*) and astrocytes (e.g. *S100A10, CD99, ID2, CRYAB, and CA2*). Intriguingly, some of those OL or astrocyte specific genes (e.g. *KCTD12, HMGCS1, FAM131C APCDD1, S100A10, CD99, and ID2*) are also expressed in primary microglia, neurons, or endothelial cells^3,34^. It needs to be considered that our *in vitro* differentiated cells may be somewhat different from their endogenous *in vivo* counterparts. On the other hand, the primary cells for the previous studies were immunopanned from a whole brain, and it is likely that OLs and astrocytes cells contaminated the microglia or endothelial populations during immunopanning. Distinction of such contaminating cells is now possible with scRNA-seq methods.

In addition to regulatory genes and TFs, miRNAs have also been implicated in OPC specification and differentiation^47–49^. Although the DropSeq approach we used does not capture mature miRNAs because they are not poly-adenylated, DropSeq can sometimes capture pri-miRNAs, the poly-adenylated precursor transcripts that give rise to miRNAS after processing by Drosha. Demonstrating the power of this approach, one of the major differentially expressed, OL enriched, pri-miRNAs that we identified is miR219-A2, which is highly enriched in human OLs^5,47^ and has previously been shown to be important for myelination and re-myelination in mice^48,50,51^. Our PD-TT reporter system could thus be a useful resource for future studies to explore ALC and OLLC specific miRNAs.

Additionally, the utility of our scRNA-seq data is further supported by IPA analysis, which revealed both known and novel pathways associated with OL and astrocyte differentiation. Of particular interest is the finding of enrichment of the CXCR4, Sphingosine-1-phosphate, integrin, mTOR and cholesterol biosynthesis signaling pathways in mature OL cells. By pharmacologically intervening these pathways, we show that targeted inhibition of *CYP51A1, TM7SF2*, and *EBP* within the Cholesterol biosynthesis pathway enhances OL differentiation, and that inhibition of mTOR signaling reduces OL differentiation in our hOPCs. Inhibition of *CYP51A1, TM7SF2*, and *EBP* gene functions has been shown to cause accumulation of 8,9-unsaturated sterols^16,18^. Since these compounds increase the amount of MBP mRNA, it is possible that the 8,9-unsaturated sterols target upstream of the MBP to increase its production. Transcriptomic study of the human OPCs treated with these compounds or supplemented with the 8,9-unsaturated sterols would help identify the upstream regulators of the MBP expression. Our pathways enrichment analysis also suggests CXCR4, Sphingosine-1-phosphate, integrin, and EIF2 as other potential targets for future OL maturation and remyelination studies. Although the CXCR4 and the S1P inhibitors we tested did not have any significant effect on the human OPCs, further studies using more specific and potent drugs that modulate these pathways are warranted.

The long-term *PDGFRα* expression in OLLCs and the heterogeneity of our PDGFR*α*+ OPCs allowed us to perform pseudotime analysis and study their differentiation trajectories. The pseudotime analysis revealed OLs vs astrocytes as the two major lineage trajectories of hOPCs, which contrasts with a recent report that suggests that Pdgfr *a+* mouse OPCs can give rise to OLs, neurons or VLMC-pericytes^23^. Furthermore, our analysis, although keeping in mind that it is an *in vitro* study with hESC-derived *PDGFRα*+ cells, suggests that astrocytes can originate not only from OPCs but also from later stage oligodendrocyte lineage cells. However, emergence of oligodendrocytes from later stage astrocyte lineage cells seems unlikely (Figure 6). We also identified transcription factors (TFs) that potentially help modulate lineage specification, differentiation and maturation of OPCs into either OLs or astrocytes. Continued upregulation of the TFs *ZEB2, TSC22D4, ARID4B, PARP1, E2F3, SOX10, TCF7L2, TSC22D1, RBPJ, ARHGAP35, SOX4, SOX11, MLLT11, RBM22, ZNF711, EZH2* and *DACH2*, which are enriched in OLs and downregulated in astrocyte cells, and *HES1, EGR1, FOSB, NFIA, NR2F1, ID3, KLF6* and *ZFP36L1*, whose expression is enriched in astrocytes and decreased in OLs, seems to drive specification of OLs vs astrocytes from PDGFR*α*+ OPCs. The role of the majority of these TFs in OL/astrocyte differentiation and maturation has not been studied. Since the function of *SOX10* and *TCF7L2* in OL development and *NFIA* in astrocyte differentiation is well known, and a crucial role of *ZFP36L1* in OL-astrocyte lineage transition was recently reported^52^, it is conceivable that the TFs we identified may play important role in OL vs astrocyte lineage specification. Loss of function and gain of function studies of these genes and TFs in OPCs will help to further validate and confirm their role in lineage specification of human OPCs.

Although mouse and human OPCs share transcriptomic similarity and conserved pathways, there appears to be some important species-specific distinctions. One example, as described above, is that human PDGFR*α*+ OPCs have the potential to become OLLCs or ALCs while the mouse PDGFR*α*+ OPCs seem to give rise primarily to OLLCs, neurons or pericytes (Figures 4 and 6)^7,9,23,53^. In addition, unlike in animal models of MS where myelin is regenerated by newly formed oligodendrocytes, the capacity to generate oligodendrocytes around the lesions of human patients is generally diminished, and the limited remyelination that does occur at MS lesions is likely generated by pre-existing OLs^54^. Thus, both for improved disease modeling and to support drug discovery efforts, it is crucial to better define the similarities and differences between murine and human myelination. We hope that the system that we described in this manuscript along with our scRNAseq dataset (http://zacklab.org/OPCs/) will help provide the basis for ongoing and future studies that will more fully define the molecular mechanisms of human myelination and re-myelination.

## ACKNOWLEDGEMENTS

We thank Valentina Fossati (The New York Stem Cell Foundation Research Institute) for helping establish the OPC differentiation protocol in the lab and James Goldman (Columbia University) for providing the O4 antibody and O4 hybridoma. This work was supported by grants from the Maryland Stem Cell Research Fund, Race to Erase MS, and NIH (P30 EY001765 and K99 EY 029011), Research to Prevent Blindness, and generous gifts from the Guerrieri Family Foundation.

## AUTHOR CONTRIBUTIONS

X.C. designed the study, conducted experiments and wrote the manuscript. A.K. performed the Dropseq experiments, analyzed RNA-seq data, and edited the manuscript. C.B assisted with experimental designs and edited the manuscript. P.D made the website. J.L.M performed the pathway enrichment analysis. C.C. and H-Q.M. provided nanofibers and assisted with the myelination assay. A.K., L.C., and T-H.W. established the Dropseq platform in the lab. P.A.C helped with experimental design and editing the manuscript. D.J.Z designed the study, edited the manuscript, and provided funding support.

## METHODS

### Human ES cells and culture conditions

hESC line WA09 (WiCell), an NIH-approved hESC line, was used for this study. hESCs were maintained in StemFlex media (A3349401, Thermo Fisher Scientific) on growth factor-reduced Matrigel (354230, Corning) coated plates at 10% CO2/5% O_2_. However, during reporter cell-line generation, the hESCs were maintained in mTeSR1 media (Stemcell Technologies). hESC colonies were passaged by dissociating with Accutase (A6964, Sigma-Aldrich). Cells were maintained in stem cell media containing 5 mM blebbistatin (B0560, Sigma-Aldrich) for the first 24 hours after passaging, to improve single cell survival. hESCs as well as purified OPCs were cryopreserved using CryoStor CS10 (07930, Stemcell Technologies) following the manufacturer’s instructions.

### Cloning

A guide sequence targeting the stop codon of the *PDGFRα* locus was cloned into the BbsI restriction site of the Cas9 plasmid (Cas9-P2A-Puro modified from Addgene #62988^29^). pSpCas9(BB)-2A-Puro (PX459) V2.0 was a gift from Feng Zhang (Addgene plasmid # 62988; http://n2t.net/addgene:62988). To clone the donor plasmid, a ~2 kb PCR product was amplified from genomic DNA extracted from H9 ES cells and cloned into Zero Blunt TOPO cloning vector (ThermoFisher Scientific). The tdTomato-P2A-Thy1.2 reporter DNA sequence was then introduced into the TOPO-based donor plasmid, precisely upstream of the *PDGFRα* stop codon using Gibson assembly (New England Biolabs).

### Generation of PDGFRα Reporter Cell Line

Gene editing and reporter cell line generation was performed using a transient antibiotic selection method^29^. Cells were transfected using the Lipofectamine Stem (STEM00001, ThermoFisher Scientific) transfection reagent following the manufacture’s recommended protocol. 0.35 μg Cas9 plasmid (Cas9-P2A-Puro modified from Addgene #62988) containing a gRNA sequence targeting the stop codon of the PDGFRα locus and 0.75 μg of donor plasmid were used for transfection. Roughly 40 hours after transfection, the cells were selected with 0.6 μg/mL of puromycin for 24 hours. Selected cells were passaged at 500–1000 single cells per well of a 6 well plate for colony formation followed by colony picking and PCR analysis^29^. PCR was performed using the Phusion Flash mastermix (ThermoFisher Scientific) and a 2-step PCR protocol following the manufacturer’s instruction.

### Oligodendrocyte Differentiation Protocol

hESCs were differentiated into OPCs and OLs as previously described ^43^ with minor modifications. Briefly, hESCs were dissociated to single cells and plated on Matrigel coated plate at 100,000 cells/well of a 6-well plate and maintained in StemFlex media at 10% CO2/ 5% O2. Two days after passaging, neural differentiation and spinal cord patterning was induced through dual SMAD inhibition and addition of 100 nM all-trans RA^31^. From day 8 to day 12, differentiating cells were maintained in neural induction media supplemented with RA (100nM) and SAG (1 mM). At day 12, adherent cells were lifted and cultured in low-attachment plates to favor sphere aggregation. At day 30, spheres were plated into poly-L-ornithine/laminin-coated dishes in a media supplemented with, B27 (Thermo Fisher, 12587010), N2 supplement (Thermo Fisher, 17502048), PDGF-AA, neurotrophin-3, HGF, and T3. Once a significant number of tdTomato+ OPCs were visible around day 65-70, mitogen free glial medium was used to drive oligodendrocyte maturation.

### Flow Cytometry and MACS Purification of the Reporter hOPCs

Flow cytometry analysis and MACS purification were performed as previously described^27^ with the following modifications. Cells were dissociated into single cell suspension by incubating in accutase for ~45 minutes. The single cell suspension was then passed through a ~70 uM cell strainer (BD Biosciences), washed, and resuspended in Live Cell Imaging Solution (ThermoFisher Scientific) for analysis with an SH-800 Cell Sorter (Sony Biotechnology, San Jose, CA). For MACS purification, cells were resuspended in MACS buffer after passing through cell strainer. A CD90.2 (THY1.2), O4 or A2B5 MicroBeads were added to the cell suspension and incubated at room temperature for 15 minutes for cell binding. Cells were generally run through the MS column twice without additional supplementation of MicroBeads to increase the purity and achieve ~90% tdTomato+ cells. All MACS reagents were purchased from Miltenyi Biotec (Auburn, CA) and manufacturer instructions were followed.

### Testing pharmacological compounds on hOPCs

Day 85 hOPCs were MACS purified to ~90% purity. 200,000 purified hOPCs were plated per well of a PLO/laminin coated 24 well tissue culture plate, in a mitogen free glial media. Day after plating the cells, culture media was replaced with media containing different compounds or DMSO. Based on previous publications and our lab’s unpublished screening work, following optimal dose for each compound was chosen: **a.** CYM5520 (1.2 uM) **b.** WZ811 (1uM) **c.** Ketoconazole (370 nM) **d.** Amorolfine (370 nM) **e.** Tasin-1 (41nM) **f.** Rapamycin (123 nM). All the compounds were purchased from Selleckchem (selleckchem.com). Media containing the compounds was replaced on day 3, and on day 7 the cells were either lysed for RNA extraction and qRT-PCR or fixed with 4% paraformaldehyde for immunostaining.

### Immunofluorescence Staining, Microscopy, and qRT-PCR

All sequences for qRT-PCR primers can be found in Table S1. Total RNA was isolated using the RNeasy Mini Kit (QIAGEN) and reverse transcribed using the High Capacity cDNA Reverse Transcription Kit (Applied Biosystems). A 2uL PCR reaction was setup using acoustic liquid handler (ECHO 550, Labcyte) and performed with the CFX384 real-time PCR instrument (Bio-Rad). All assays included technical and biological triplicates were analyzed using the Sso Advanced Universal SYBR Green Supermix (Bio-Rad).

For immunofluorescence staining, cells were fixed with 4% paraformaldehyde, and simultaneously permeabilized and blocked with 0.2% Triton X-100 + 5% BSA + 5% normal goat serum (or a serum specific to the host of secondary antibody) for an hour. Cells were then incubated with appropriate dilution of primary antibodies over-night followed by secondary antibodies for 2 hours. Fluorescence images were taken using either the EVOS FL Auto 2 (ThermoFisher Scientific) or Zeiss 510 confocal microscope.

#### Live cell imaging

The EVOS FL Auto 2 Cell Imaging System (ThermoFisher Scientific) was used for imaging cells in culture over time for time-lapse videos. Cells were maintained in a live cell chamber at 37°C with 5% CO2 and 85% humidity and areas around neurospheres were scanned every 20 minutes. Images were compiled at 15 fps using ImageJ to generate timelapse videos.

### Drop-seq-based Single Cell Capture and RNA-sequencing

Drop-seq-based single cell RNAseq was performed as previously described by Macosko et al.^14^. Barcoded microparticles were purchased from Chemgenes Corporations. Using the microfluidic device, single MACS purified PD-TT reporter cells taken at 77, 89 and 104 days were captured into a ~1 nL size droplets containing barcoded nanoparticles and lysis buffer. Generated droplets were broken with perfluorooctanol (Sigma, 370533) in 30 ml of 6× SSC. The beads were then washed, reverse transcribed, PCR amplified, and the amplified cDNA quantified using a BioAnalyzer High Sensitivity Chip (Agilent). The cDNA was then fragmented and amplified for 3’ prime end sequencing with the Nextera XT DNA sample prep kit (Illumina). The libraries were purified, quantified, and then sequenced on the rapid flow chip in Illumina HiSeq 2500.

### Statistical Analysis

All qRT-PCR data are presented as fold change in RNA normalized to *GAPDH* expression. qRT-PCR data was analyzed using qbase+ qPCR analysis software and graphed using Prism (GraphPad). Data are presented as +/- SEM unless stated otherwise.

### Bioinformatic Analysis

The principal component analysis (PCA) and t-distributed stochastic neighbor embedding (tSNE) analyses were performed using a previously published R package, Seurat^33^. As a quality control, only cells that had a minimum of 250 mRNA molecules and a maximum of 20% mitochondrial RNA were used for analysis. Cells with greater than 30,000 UMIs were also filtered out to remove probable doublets. Genes that were expressed in a minimum of 3 cells were included for the analysis. 1874 highly variable genes were input for PCA analysis, and the 16 statistically significant PC’s were used for clustering and t-SNE analysis. Cluster identity was called based on previously published marker genes. Time-series analysis to generate a pseudotemporal trajectory was performed using an unsupervised differential gene expression test based on sample age in Monocle, following previously published detailed instructions ^24,55^. The top 752 genes differentially expressed based on age were used for ordering and trajectory reconstruction. Differential gene expression was performed on each node of the resulting trajectory to identify genes with branch-dependent expression. Differential gene expression was performed using either Seurat or Monocle. To calculate Spearman Correlation, 767 highly variable genes from each human tissue-type published by Zhang *et al.^34^* was compared to each of our clusters. For comparison with mouse cells, all the genes expressed in each of our clusters were compared to the genes from each of the mouse CNS cell-type previously published^3^.

The networks and pathway analyses were generated through the use of IPA (QIAGEN Inc., https://www.qiagenbio-informatics.com/products/ingenuity-pathway-analysis)^56^. For the analysis, differentially expressed genes and their corresponding p-value and fold change (from supplemental table 1) was uploaded for each cluster. The sterol biosynthesis pathway map was generated using KEGG via Visualization and Integrated Discovery (DAVID) tools and its steroid biosynthesis pathway as the reference pathway.

## DATA AND SOFTWARE AVAILABILITY

RNA-Seq data generated for this paper will be deposited to the NCBI’s Gene Expression Omnibus (GEO) database (https://www.ncbi.nlm.nih.gov/geo/), and will also be available for viewing and download at http://zacklab.org/OPCs/.

## Supplemental Figures and Figure Legends

**Supplemental Figure 1.**
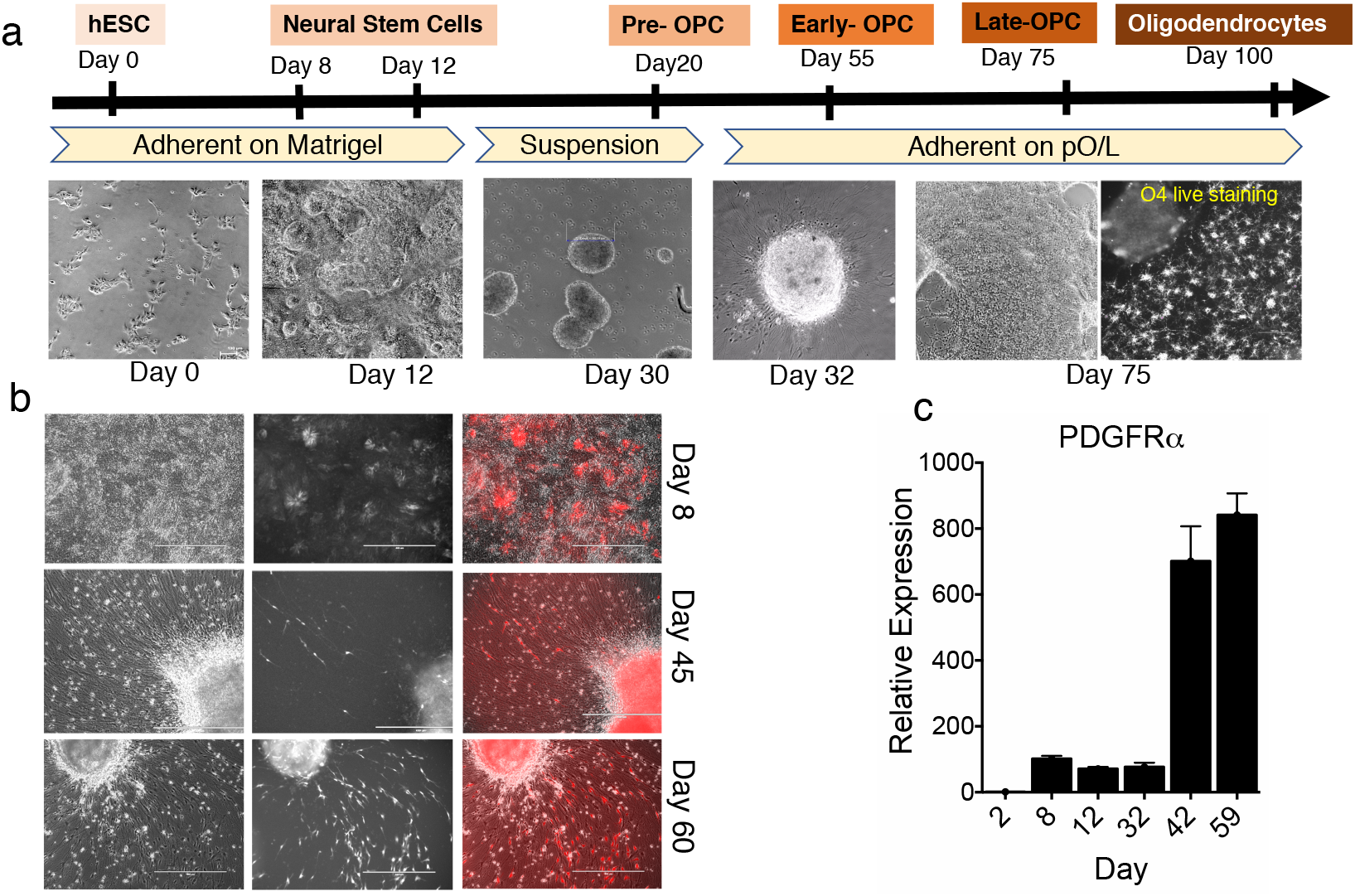
Differentiation of PDGFR*∝* reporter (PD-TT) cell line. **a)** Timeline of the differentiation protocol with images at different stages of differentiation. At Day 75, the majority of the OPCs that migrate out from the neurospheres plated on day 30 are O4 positive. **b)** tdTomato expression (driven by PDGFRα) is observed as early as day 8 on the neuroepithelial layer of the differentiating culture. However, individual, tdTomato+/PDGFRα positive cells emerge from the neurospheres around day 40 and rapidly divide and migrate out as seen in day 60 culture. **c)** qPCR analysis of the *PDGFRα* mRNA level coincides with the tdTomato expression in the cultures.

**Supplemental Figure 2.**
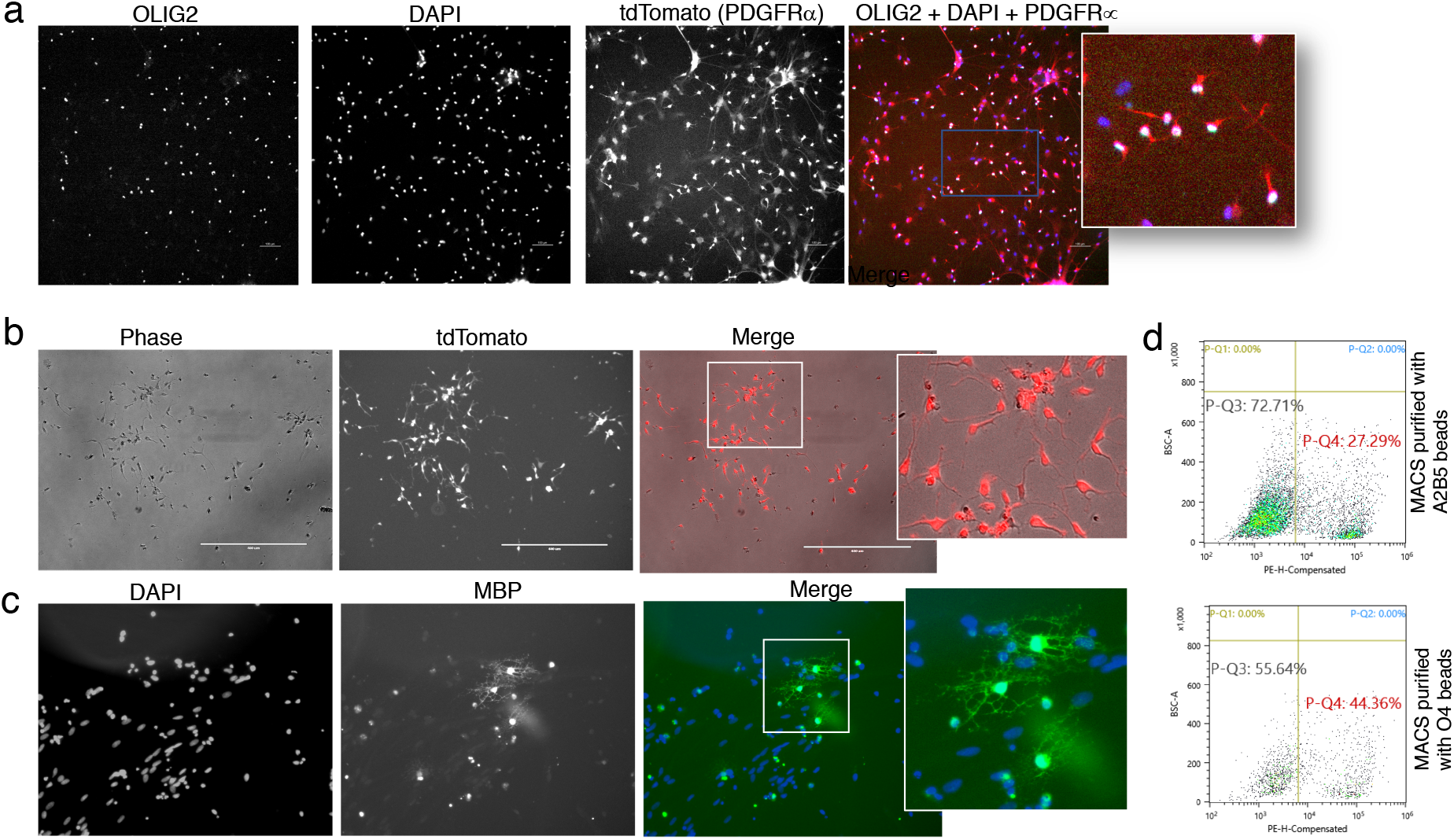
Reporter OPCs maintain competence after freezing. **a)** Immunostaining to show purified tdTomato+/PDGFRα+ cells express OLIG2. **b)** MACS purified and cryopreserved tdTomato+/PDGFRα+ cells plated on PLO-laminin coated plates for two days. **c)** Cryopreserved cells mature into MBP+ OLs within three weeks in mitogen-free glial media. **d)** PD-TT cells at day 77 of differentiation enriched using A2B5 or O4 microbeads were FACS analyzed for the expression of tdTomato. ~27% of A2B5+ and ~44% of O4+ cells also express tdTomato/PDGFR*α*.

**Supplemental Figure 3.**
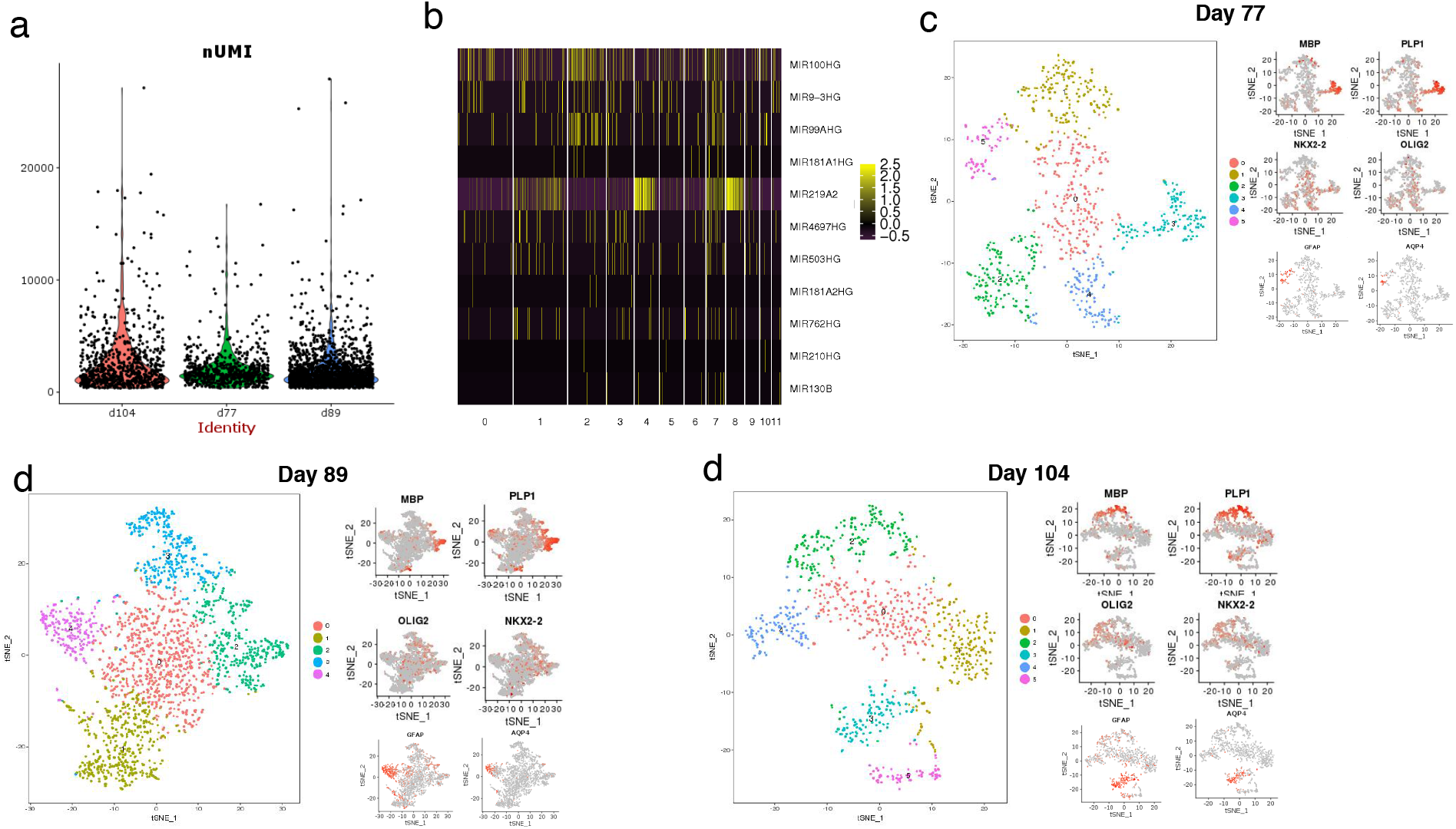
Bioinformatic analysis of scRNAseq data. **a)** Violin plot showing number of UMIs by age. In order to remove probable doublets, cells with UMI above 30,000 were removed from the analysis. **b)** Heatmap of pri-microRNAs captured by scRNAseq analysis. miR219 is enriched in OL clusters, miR99AHG and miR100HG are enriched in astrocyte clusters. **c-e)** t-SNE based unsupervised clustering of single cells captured from PDGFRα+/tdTomato+ reporter OPCs at different stages of differentiation (day 77, day 89, and day 104). Cluster specific enrichment of various OL and astrocyte markers indicate OLLC and ALC subgroups of PDGFRα+ cells in each time-point.

**Supplemental Figure 4.**
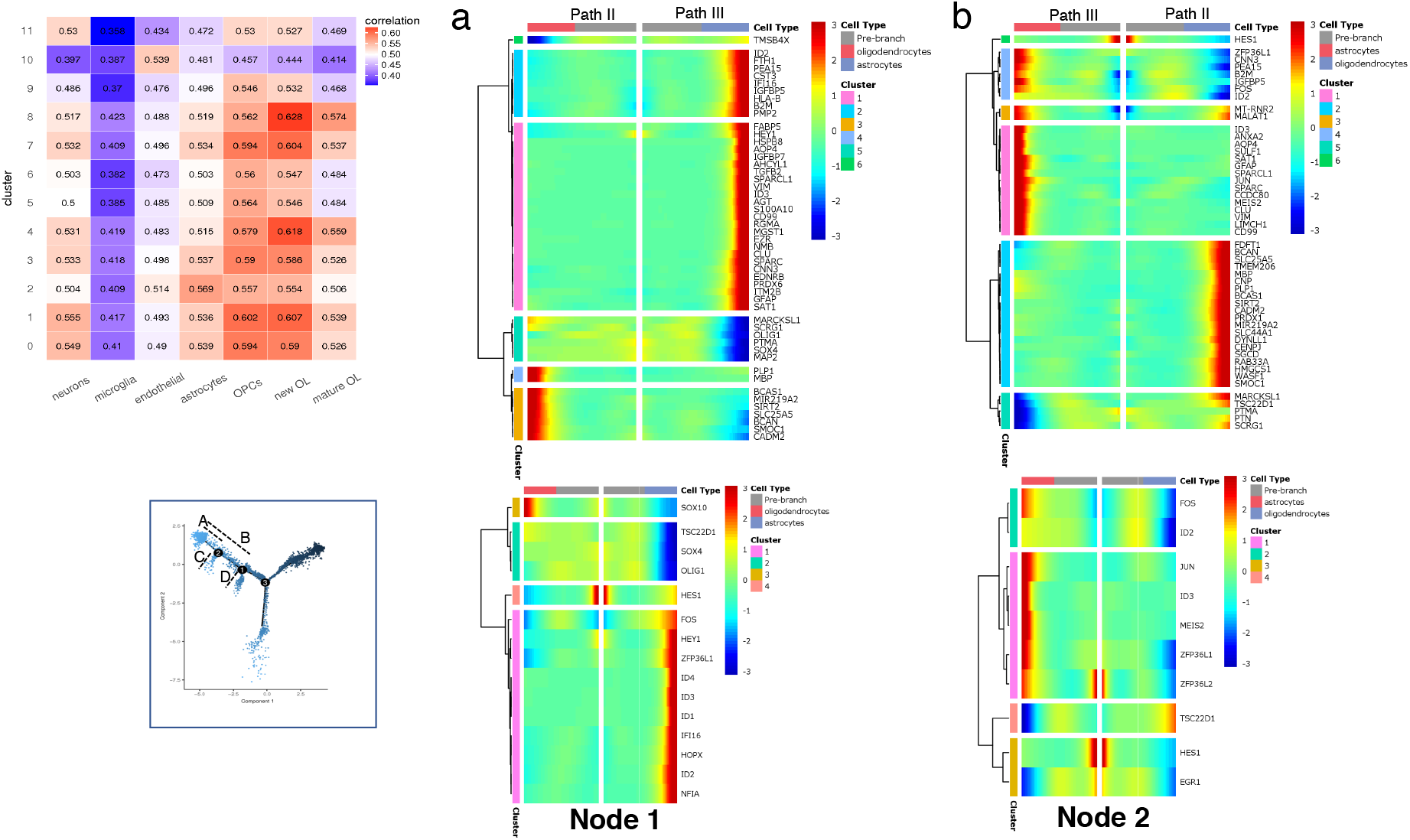
Heatmap of the most differentially expressed genes and transcription factors between different sub-branches. **a)** Spearman correlation of single cell transcriptome-based clusters with RNAseq data from primary mouse brain cells^3^. **b and c)** Analysis of differential expression at node 2 represents genes from sub-branch A vs C **(b)**, and node 1 represents sub-branch B vs D **(c**). See inset on the lower left for labelling of nodes and sub-branches.

**Supplemental Figure 5.**
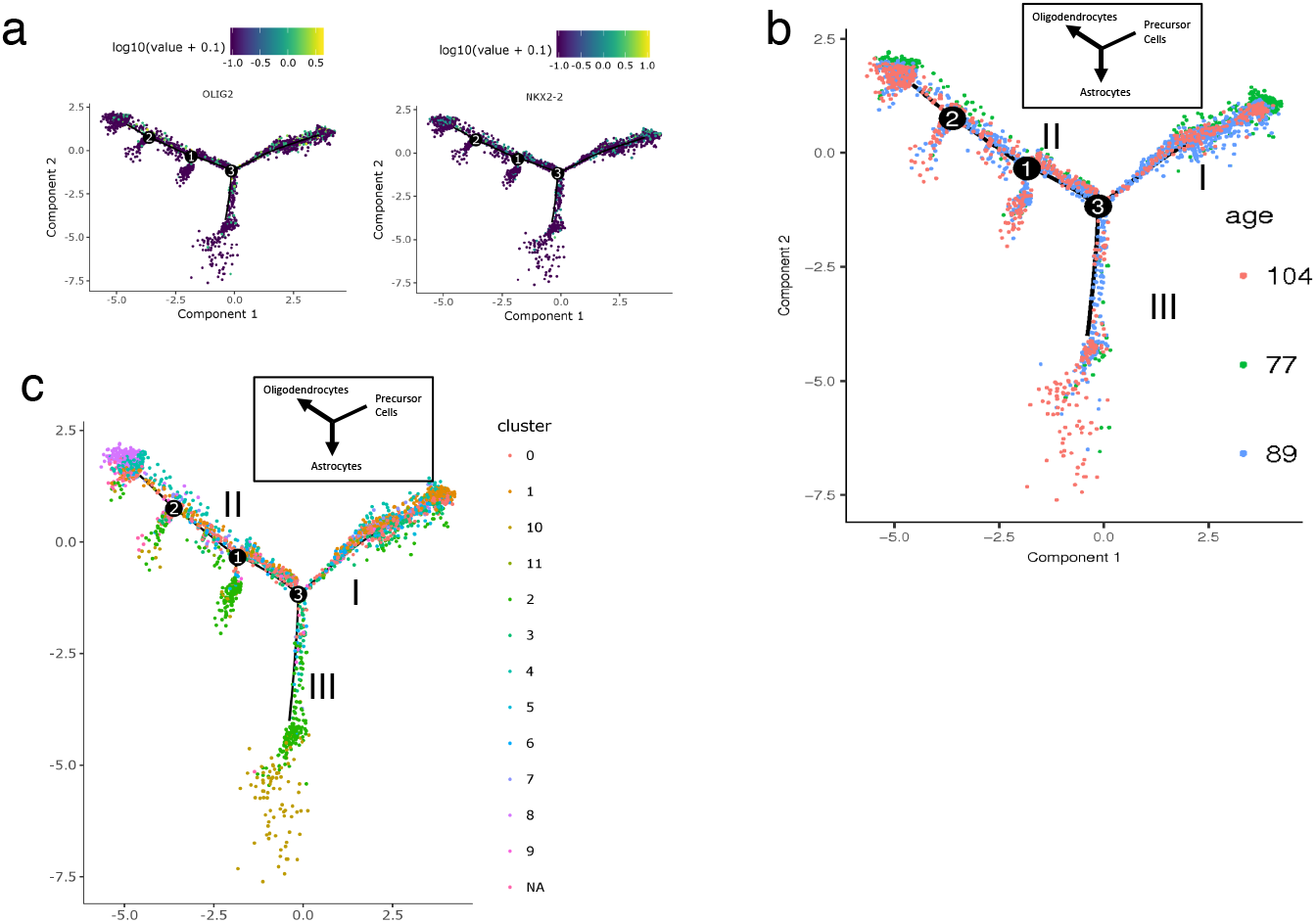
**a)** Expression of OPC markers within the trajectories show higher expression in Path I and II, but not in Path III or the smaller branches that emerge along Path II. **b)** Cells in each path of the trajectory are labelled by day at which they were purified (day 77, 89, or 104). Each path consists of cells from all three ages. **c)** Overlay of cells from each cluster from figure 4a onto the pseudo-temporal trajectory showing where in the trajectory the cells from each cluster are located. Cells from cluster 0, 1, and 3 make up the majority of Path I population. Cells from astrocyte cluster (cluster 2) make up the majority of population on Path III (state 2 cells) and the sub-branches within Path II (state 5 and 7 cells). Cells from cluster 4 and 8 are located at the end of Path II (state 6 cells). Cells from cluster 10 are located at the end of Path III or state 2 cells.

**Supplemental Video 1.** A 6 hour long time-lapse video of the PD-TT reporter cells at day 45 of differentiation. First few tdTomato+ OPCs are seen migrating out of the neurosphere (bottom of the image) plated on a surface coated with poly-L-ornithine and laminin.

**Supplemental video 2.** Time-lapse video of the PD-TT reporter cells at day 65 of differentiation taken over 18 hours. Numerous tdTomato+ OPCs have migrated out of the neurosphere (at the center of the image) and are seen moving around on a coverslip coated with poly-L-ornithine and laminin.

**Supplemental Table 1**. Oligonucleotides used for this study.

**Supplemental Table 2.** List of the differentially expressed genes from each cluster.

**Supplemental Table 3**. List of the Ingenuity canonical pathways and their associated transcripts for each cluster.

